# Rapid bacterial identification by direct PCR amplification of 16S rRNA genes using the MinION^TM^ nanopore sequencer

**DOI:** 10.1101/435859

**Authors:** Shinichi Kai, Yoshiyuki Matsuo, So Nakagawa, Kirill Kryukov, Shino Matsukawa, Hiromasa Tanaka, Teppei Iwai, Tadashi Imanishi, Kiichi Hirota

## Abstract

Rapid identification of bacterial pathogens is crucial for appropriate and adequate antibiotic treatment, which significantly improves patient outcomes. 16S ribosomal RNA (rRNA) gene amplicon sequencing has proven to be a powerful strategy for diagnosing bacterial infections. We have recently established a sequencing method and bioinformatics pipeline for 16S rRNA gene analysis utilizing the Oxford Nanopore Technologies MinION™ sequencer. In combination with our taxonomy annotation analysis pipeline, the system enabled the molecular detection of bacterial DNA in a reasonable timeframe for diagnostic purposes. However, purification of bacterial DNA from specimens remains a rate-limiting step in the workflow. To further accelerate the process of sample preparation, we adopted a direct PCR strategy that amplifies 16S rRNA genes from bacterial cell suspensions without DNA purification. Our results indicate that differences in cell wall morphology significantly affect direct PCR efficiency and sequencing data. Notably, mechanical cell disruption preceding direct PCR was indispensable for obtaining an accurate representation of the specimen bacterial composition. Furthermore, 16S rRNA gene analysis of mock polymicrobial samples indicated that primer sequence optimization is required to avoid preferential detection of particular taxa and to cover a broad range of bacterial species. This study establishes a relatively simple workflow for rapid bacterial identification via MinION^TM^ sequencing, which reduces the turnaround time from sample to result, and provides a reliable method that may be applicable to clinical settings.

## Introduction

Acute infectious diseases remain one of the major causes of life-threatening conditions with high mortality, particularly in patients under intensive care. Therefore, rapid and accurate identification of pathogenic bacteria facilitates the initiation of appropriate and adequate antibiotic treatment [1, 2]. Although culture-based techniques are still the forefront of clinical microbial detection, these methods are time-consuming and have the critical drawback of not being applicable to non-cultivable bacteria [3].

As an alternative approach for overcoming the limitations of traditional culture-based bacterial identification, metagenomic sequencing analysis has been introduced for the diagnosis of bacterial infections [4]. Among the sequence-based microbiome studies, the 16S ribosomal RNA (rRNA) genes have been the most predominantly used molecular marker for bacterial classification [5]. The bacterial 16S rRNA gene is approximately 1,500 bp long and contains both conserved and variable regions that evolve at different rates. The slow evolution rates of the former regions enable the design of universal primers that amplify genes across different taxa, whereas fast-evolving regions reflect differences between species and are useful for taxonomic classification [6].

Targeted amplification of specific regions of the 16S rRNA gene followed by next generation sequencing (NGS) is a powerful strategy for identifying bacteria in a given sample. Despite the high-throughput capacity, second generation DNA sequencing technologies provide relatively short read lengths with limited sequence information, which often hampers accurate classification of the bacterial species [7]. A portable sequencing device MinION™ from Oxford Nanopore Technologies offers a number of advantages over existing NGS platforms [8, 9]. Besides its small size and low cost, the intriguing feature of MinION sequencer is that it can provide a real-time and on-site analysis of any genetic material, which should be useful especially for clinical applications [10]. With the ability to generate longer read lengths, MinION™ analysis targets the whole coding region of the 16S rRNA gene, showing great potential for rapid pathogen detection with more accuracy and sensitivity [11-17]. We have previously established a sequencing method and bioinformatics pipeline for rapid determination of bacterial composition based on 16S rRNA gene amplicon sequencing via the MinION™ platform [15]. A 5-minute data acquisition using MinION™ and sequence annotation against our in-house genome database enabled the molecular detection of bacterial DNA in a reasonable timeframe for diagnostic purposes.

In the current study, we attempted to further refine and update the protocols for 16S rRNA gene sequencing analysis. We evaluated the performance of primer sets targeting the near full-length 16S rRNA gene. To accelerate the process of sample preparation, we adopted a direct polymerase chain reaction (PCR) strategy to amplify the 16S rRNA gene from bacterial extracts without DNA purification.

## Materials and Methods

### Direct PCR amplification of 16S rRNA genes

The number of colony forming units (CFU) of bacteria (*Escherichia coli* and *Staphylococcus aureus*) was determined by plating serial dilutions of cultures on agar plates and counting colonies [18]. For mechanical cell disruption, zirconia beads (EZ-Beads™; Promega, Madison, WI) were added to the bacterial cell suspensions and the samples were vortexed for 30 s. The bacterial cell samples with or without mechanical disruption were added directly to PCRs for amplifying the 16S rRNA genes. Bacterial DNA was purified using DNeasy Blood & Tissue Kit (Qiagen, Hilden, Germany) and used as a PCR template. PCR amplification of 16S rRNA genes was conducted using the 16S Barcoding Kit (SQK-RAB204; Oxford Nanopore Technologies, Oxford, UK) containing the 27F/1492R primer set [19, 20] and LongAmp™ Taq 2X Master Mix (New England Biolabs, Ipswich, MA). Amplification was performed using an Applied Biosystems Veriti™ Thermal Cycler (Thermo Fischer Scientific, Waltham, MA) with the following PCR conditions, initial denaturation at 95 °C for 3 min, 25 cycles of 95 °C for 20 s, 55 °C for 30 s, and 65 °C for 2 min, followed by a final extension at 65 °C for 5 min. To determine the effects of human DNA contamination on 16S rRNA gene amplification, genomic DNA purified from the human monocytic cell line THP-1 was mixed with *E. coli* DNA and subjected to PCR. To amplify human β-globin gene as an internal control for the human genome, the following primers were used, forward, 5’-GGTTGGCCAATCTACTCCCAGG-3’ and reverse, 5’-TGGTCTCCTTAAACCTGTCTTG-3’. Quantitative real-time PCR was performed using SYBR Green I fluorescence and Rotor-Gene Q cycler (Qiagen). Melting-curve analysis was done using Rotor-Gene Q series software version 2.1.0.

### Genomic DNA from a mock bacterial community

MSA-1000™ 10 Strain Even Mix Genomic Material was obtained from the American Type Culture Collection (ATCC; Manassas, VA). The DNA mixture (1 ng) was used as a template for amplifying 16S rRNA genes. PCR amplification was conducted using the 16S Barcoding Kit and LongAmp™ Taq 2X Master Mix following the thermal cycling protocol as described above. Alternatively, 16S rRNA genes were amplified using KAPA2G™ Robust HotStart ReadyMix PCR Kit (Kapa Biosystems, Wilmington, MA). Amplification conditions for fast PCR using the KAPA2G™ polymerase were as follows, initial denaturation at 95 °C for 3 min, 25 cycles of 95 °C for 15 s, 55 °C for 15 s, 72 °C for 30 s, followed by a final extension at 72 °C for 1 min.

### Whole cell mock bacterial community

MSA-3000™ 10 Strain Mix Whole Cell Material was obtained from ATCC. Lyophilized bacterial cell pellets were suspended in phosphate-buffered saline (PBS) and divided into aliquots. The resulting cell suspensions were then either used for direct PCR to amplify the 16S rRNA genes (2.5 × 10^4^ cells/reaction) or subjected to mechanical cell disruption via bead-beating prior to PCR amplification. Bacterial DNA purified from the cell suspension was also used for 16S rRNA amplicon sequencing.

### Sequencing of 16S rRNA gene amplicons

PCR products were purified using AMPure XP (Beckman Coulter, Indianapolis, IN) and quantified by a NanoDrop (Thermo Fischer Scientific). A total of 100 ng DNA was used for library preparation and MinION™ sequencing was performed using R9.4 flow cells (FLO-MIN106; Oxford Nanopore Technologies) according to manufacturer’s instructions. MinKNOW software ver. 1.11.5 (Oxford Nanopore Technologies) was used for data acquisition.

### Bioinformatics analysis

MinION™ sequence reads (i.e., FAST5 data) were converted into FASTQ files by using Albacore software ver. 2.2.4 (Oxford Nanopore Technologies). Then, the FASTQ files were converted to FASTA files using our own program. In these reads, simple repetitive sequences were masked using TanTan program ver. 13 with default parameters [21]. To remove reads derived from humans, we searched each read against the human genome (GRCh38) using minimap2 with default parameters [22]. Then, unmatched reads were regarded as reads derived from bacteria. For each read, a minimap2 search with 5,850 representative bacterial genome sequences stored in the GenomeSync database (http://genomesync.org) was performed. Next, we chose species showing the highest minimap2 score as the existing species in a sample. Taxa were determined using our in-house script based on the NCBI taxonomy database [23] and then visualized using Krona Chart [24]. Sequence data from this article have been deposited in the DDBJ DRA database (https://www.ddbj.nig.ac.jp/dra/index-e.html) under accession numbers DRR157203 to DRR157213.

### Statistical analysis

For permutational multivariate analysis of variance (PERMANOVA), Morisita’s index of similarity ranging from 0 (no similarity) to 1 (complete similarity in composition) was used [25]. PERMANOVA tests were performed using R package vegan [26].

## Results

### Direct PCR approach for amplifying 16S rRNA genes from crude bacterial extracts

To overcome the time-consuming and laborious process of sample preparation for DNA sequencing, we tried to amplify the 16S rRNA gene directly from bacterial suspensions without a DNA purification step (Fig. 1). We used a commercially available kit (16S Barcoding Kit; Oxford Nanopore Technologies) with primers optimized for 16S rRNA amplicon sequencing on the MinlON™ platform. The primers were designed to amplify the near full-length 16S rRNA gene for bacterial identification [19, 20]. Each indexing primer has a unique barcode for multiplexing and contains a tag sequence at the 5’-end for attachment of sequencing adapters.

**Fig. 1.**
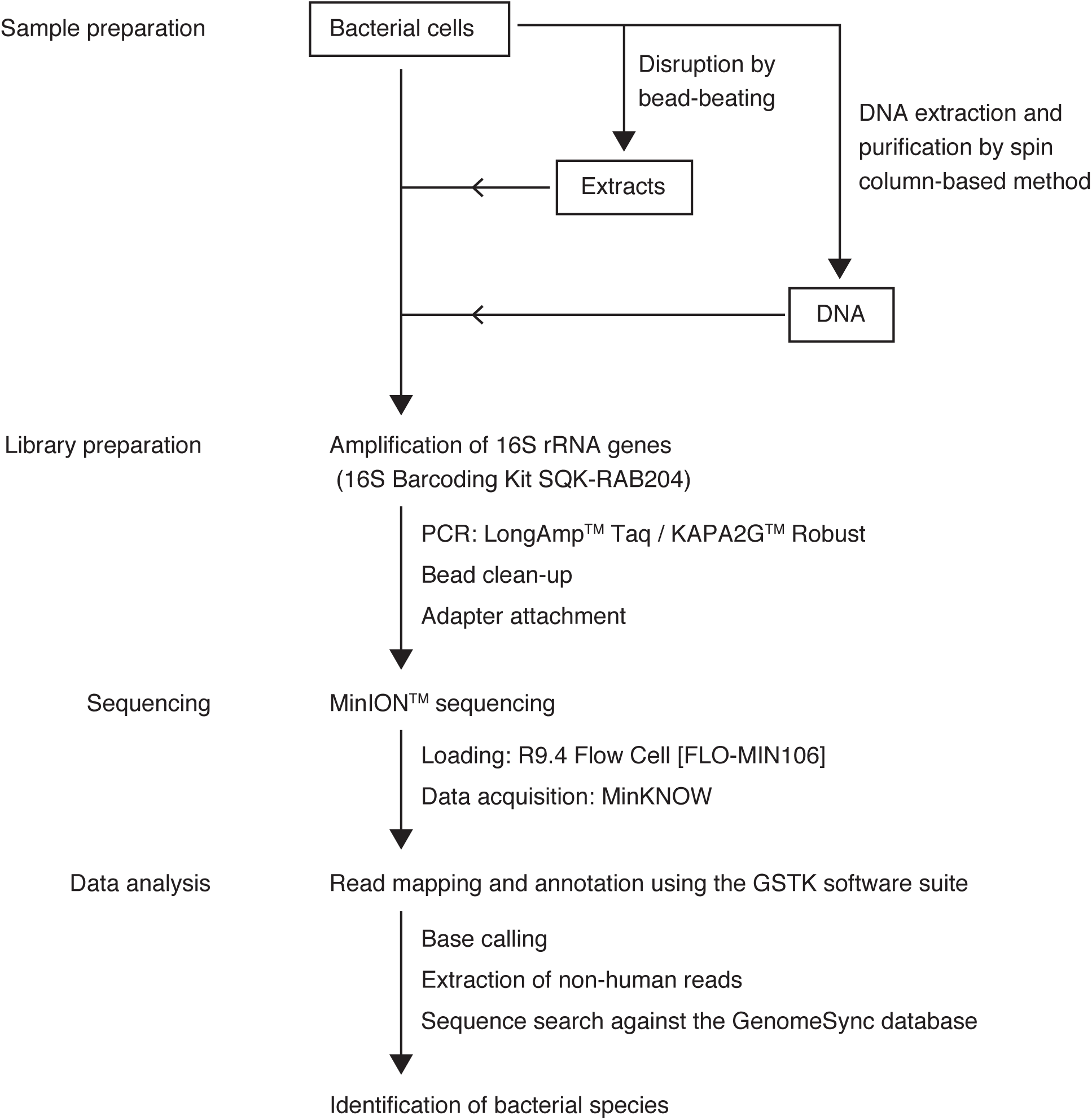
Workflow of 16S rRNA amplicon sequencing on a MinION™ platform and bioinformatics analyses. Bacterial cells were left untreated or disrupted by bead-beating and then subjected to direct PCR for amplifying the near full-length 16S rRNA genes. Additionally, purified bacteria DNA was used for 16S rRNA gene amplification. The samples were sequenced on a MinION™ platform. The data obtained were analyzed using the computational analysis pipeline Genome Search Toolkit (GSTK) with GenomeSync database.

Performance of the barcoded primers for 16S rRNA gene amplification was evaluated by PCR assays using bacterial cell suspensions. *E. coli* was chosen to represent gram-negative pathogens and *S. aureus* was used as a model for hard-to-lyse bacteria with gram-positive cell walls. A defined amount of each bacteria was serially diluted and the resulting cell suspensions were directly added to PCRs with LongAmp™ Taq DNA polymerase. As for *E. coli* suspensions, the lower limit of detection was less than 1 × 10^2^ CFU in agarose gel electrophoresis (Fig. 2A). On the other hand, a higher number of cells was required for detecting 16S rRNA gene amplicons from *S. aureus* suspensions, whose detection limit was as low as 1 × 10^3^ CFU (Fig. 2B).

**Fig. 2.**
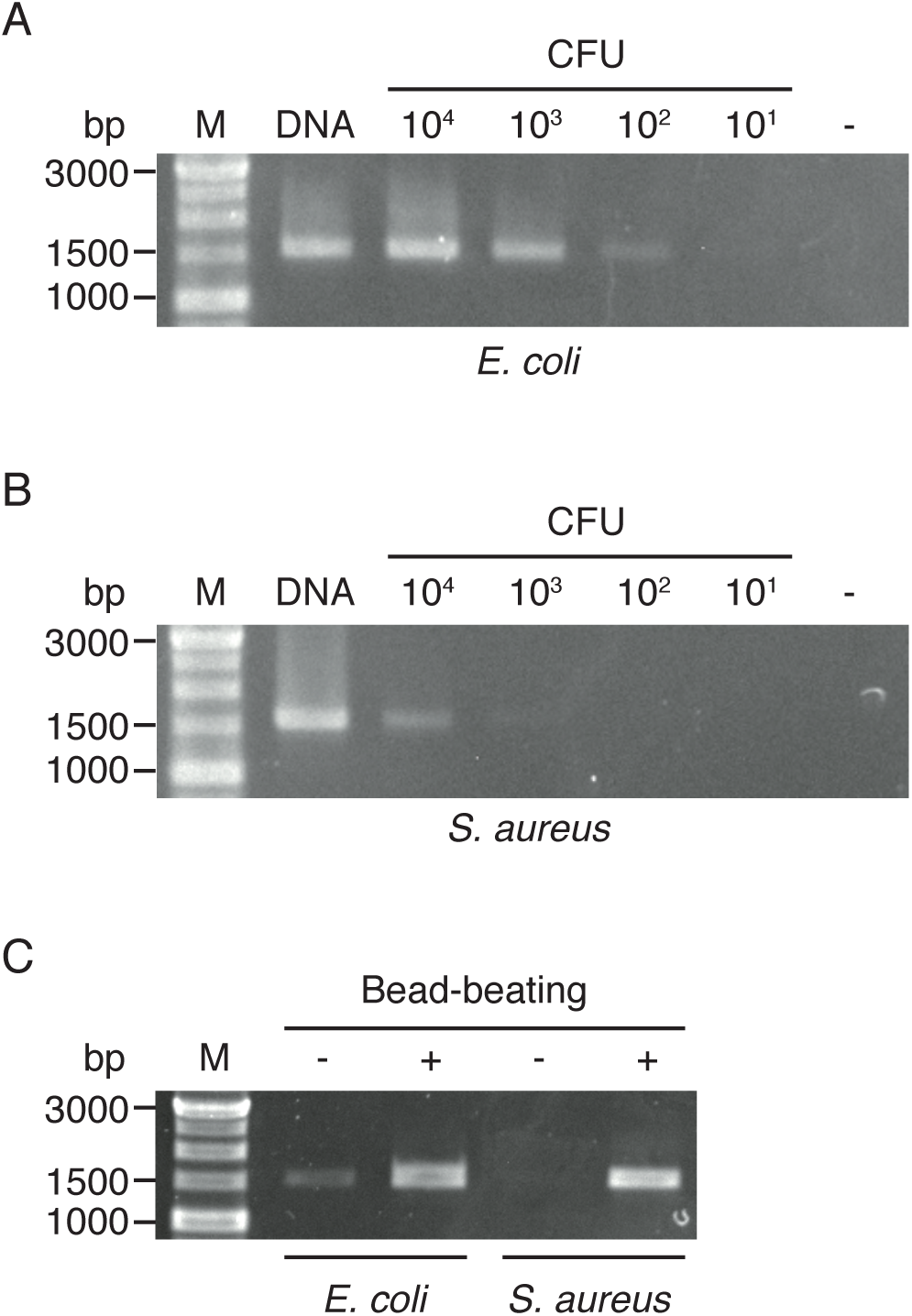
Direct PCR amplification of 16S rRNA genes without DNA purification. (A and B) A known amount of (A) *E. coli* and (B) *S. aureus* were added directly to PCRs (1 × 10^1^−1 × 10^4^ CFU/reaction) and then the amplified products were analyzed by agarose gel electrophoresis. M, molecular weight marker; DNA, 1 ng of bacterial DNA (positive control); -, no template (negative control); CFU, colony forming unit. (C) Bacterial cells were left untreated (-) or mechanically disrupted by bead-beating (+) and then the resulting cell extracts were subjected to 16S rRNA gene amplification.

To facilitate the release of bacterial DNA, cells in suspensions were disrupted by vortexing with zirconia beads before being subjected to PCR amplification. Bead-beating proved to be effective for the direct amplification of 16S rRNA genes from cell suspensions of both *E. coli* and *S. aureus*, improving the yield of PCR products (Fig. 2C).

### Rapid detection and identification of bacterial strains via direct PCR amplicon sequencing on MinION™

Having demonstrated the efficacy of the barcoded primers for amplifying 16S rRNA genes directly from bacterial suspensions, we investigated whether the direct PCR method can impact MinION™ sequencing results and the accuracy of strain identification. Bacterial cell suspensions of *E. coli* and *S. aureus* were used for preparing 16S rRNA gene amplicon libraries and then the samples were sequenced on MinION™ for 5 minutes (Table 1 and Fig. 3). In addition, sequencing libraries were prepared using purified bacterial DNA templates as a standard reference for comparison. Sequencing reads were analyzed using a bioinformatics pipeline based on a BLAST search against our in-house genome database GenomeSync. MinION™ sequencing data identified the bacteria at the species level with more than 90% of reads being correctly assigned to each species (Figs 3A and 3B). *Shigella flexneri* was additionally detected at a low abundance probably due to its high sequence similarity to *E. coli* [27, 28]. Moreover, the type of PCR template (purified DNA versus cell suspension) did not substantially affect the quality of sequence reads nor bacterial identification results (Table 1). These results demonstrate the utility of the direct PCR method, which can enable rapid pathogen identification from crude materials without the need for DNA purification.

**Table 1.**
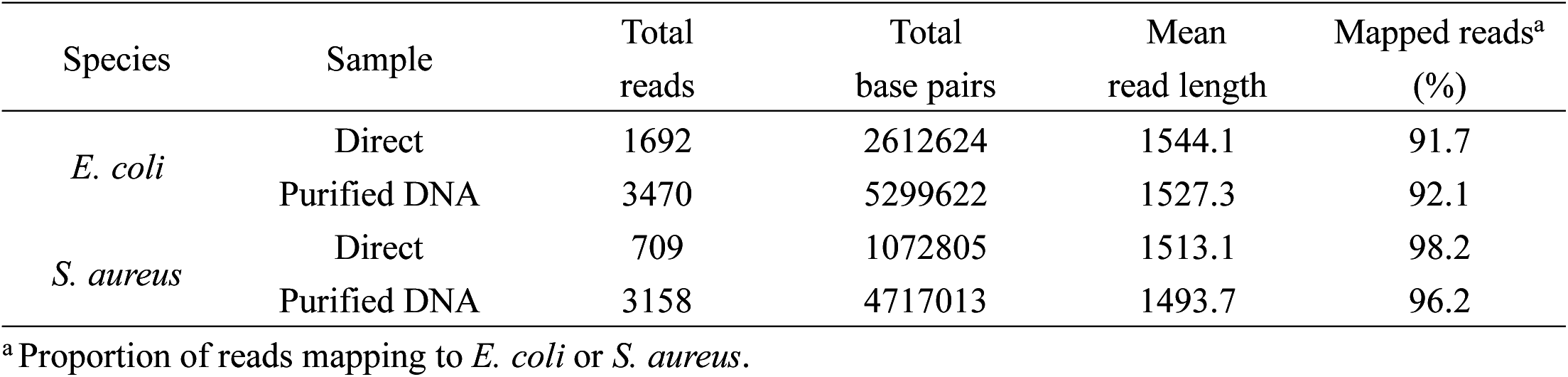
MinION sequencing of 16S rRNA gene amplicons.

**Fig. 3.**
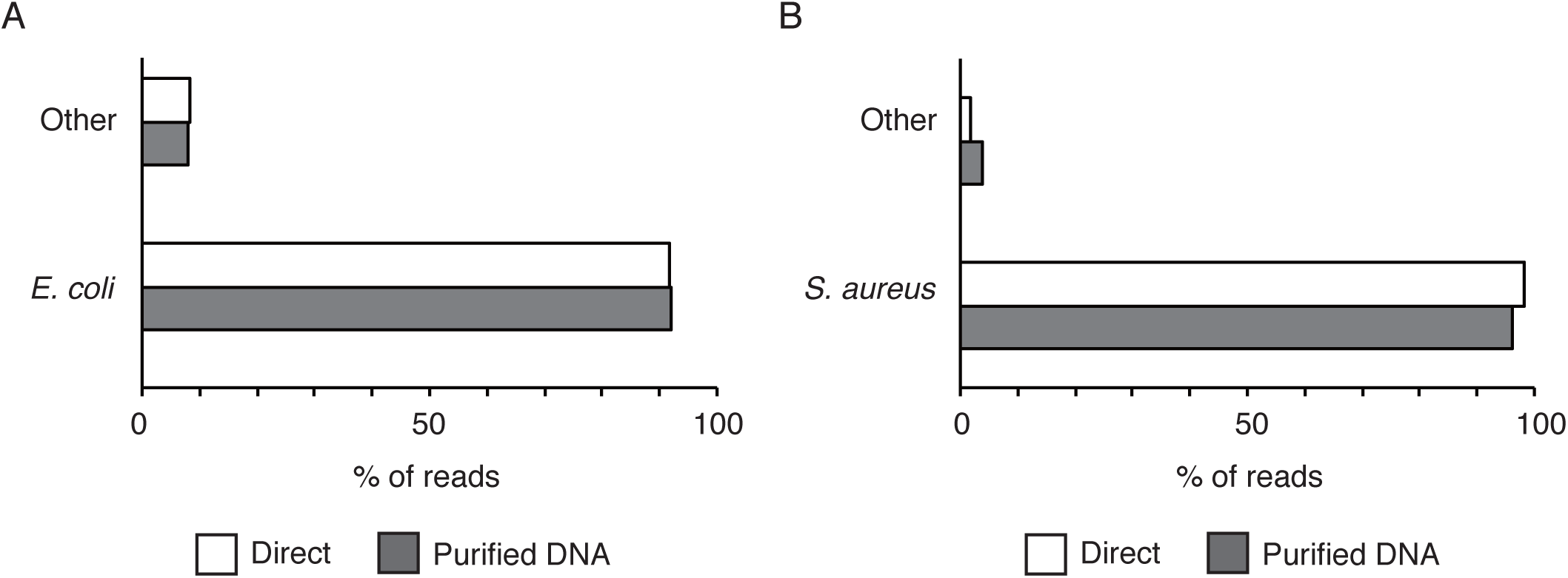
Accurate taxonomic assignment of MinION™ sequence reads amplified directly from bacterial cell suspensions. (A and B) 16S rRNA gene sequencing libraries were prepared using purified DNA templates or amplified directly from cell suspensions. The samples were sequenced on a MinION™ platform and taxonomic assignment was performed with the analytical pipeline GSTK. The classification accuracy is shown for (A) *E. coli* and (B) *S. aureus*.

### Impact of non-bacterial DNA contamination on 16S rRNA gene amplification

Successful identification of infectious pathogens should rely on the specific amplification of bacterial target sequences in clinical samples, which can often be contaminated with patient-derived human genetic materials. We tested whether a higher amount of human DNA would affect the amplification of bacterial 16S rRNA genes. *E. coli* DNA (0.1 ng) was mixed with increasing amounts of human DNA samples extracted from the monocytic cell line THP-1, after which the mixture was subjected to PCR amplification of the 16S rRNA gene (Fig. 4A; upper panel). Primers targeting the β-globin gene were used as internal control for the human genome (Fig. 4A; lower panel). Bacterial 16S rRNA genes were specifically amplified even in a background of high human DNA concentrations. The contaminated human DNA had no significant inhibitory effects on PCR product yield, which was further confirmed by quantitative real-time PCR (Fig. 4B). Melting curve analysis suggested that targeted 16S rRNA amplicons were specifically generated by PCR (Fig. 4C). Thus, the existence of non-bacterial genetic materials in the sample does not affect the sensitivity and specificity of 16S rRNA gene detection.

**Fig. 4.**
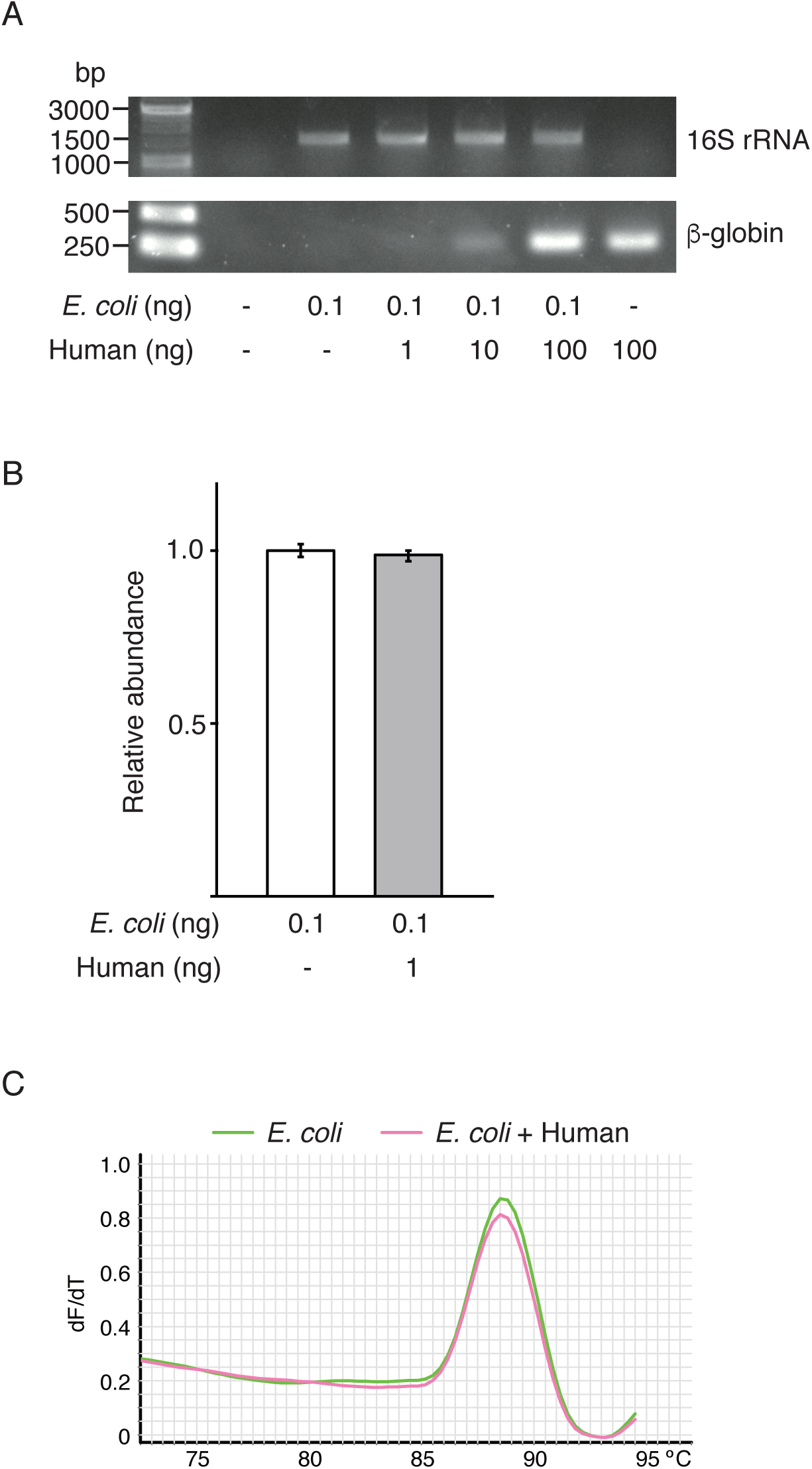
Impact of background human DNA contamination on the specific amplification of 16S rRNA genes. (A) *E. coli* DNA (0.1 ng) was mixed with varying amounts of human DNA samples and subjected to 16S rRNA gene amplification. β-globin was used as an internal control for the human genome. (B and C) *E. coli* DNA (0.1 ng) mixed with or without human DNA (1 ng) was used for quantitative real-time PCR analysis targeting the 16S rRNA gene. The graph shows the relative abundance of 16S rRNA gene amplicons. The experiment was done in duplicate. Data represent the mean values ± SD of three experiments (B). Melting curve analysis was performed with stepwise increases of 1°C (C).

### 16S rRNA gene sequencing of a mock bacterial community

The performance of the current tools for 16S rRNA gene amplicon sequencing was further tested with a mixture of DNA prepared from 10 different bacterial species. The relative abundance of individual bacterial taxa was estimated by genome size and copy number of the 16S rRNA gene (Table 2). The mock community DNA mixture was used as a template for PCR and the 16S rRNA gene amplicon libraries were sequenced on MinlON™. Nine out of 10 bacterial strains were successfully identified at the species level and PERMANOVA showed no significant community difference (*P* = 0.5) between data collected at different time points (Fig. 5). Thus, three minutes of run time generating 3,985 reads was sufficient for identifying the nine species, whereas longer run times (5 minutes, 10,167 reads; 30 minutes, 44,248 reads) did not significantly affect species detection accuracy (Table 3). There were some biases observed in the taxonomic profile; *Bacillus cereus* was detected at lower abundances than expected, while *Clostridium beijerinckii* and *E. coli* were overrepresented. These instances of partially-biased assignment or misidentification of bacterial species were not resolved by increasing the number of sequencing reads analyzed (Fig. 5). Our approach failed to identify *Bifidobacterium adolescentis* in the mock community even when it was represented in the database.

**Table 2.**
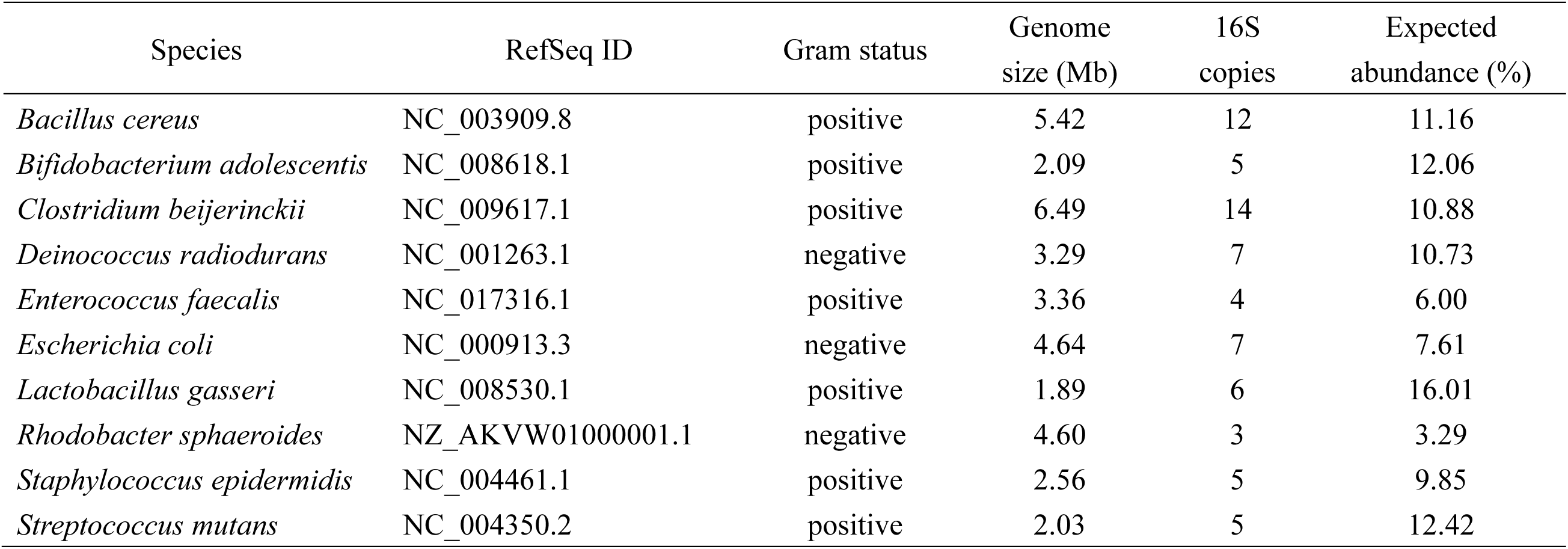
Mock community of 10 bacterial species.

| Species | RefSeq ID | Gram status | Genome size (Mb) | 16S copies | Expected abundance (%) |
| --- | --- | --- | --- | --- | --- |
| *Bacillus cereus* | NC_003909.8 | positive | 5.42 | 12 | 11.16 |
| *Bifidobacterium adolescentis* | NC_008618.1 | positive | 2.09 | 5 | 12.06 |
| *Clostridium beijerinckii* | NC_009617.1 | positive | 6.49 | 14 | 10.88 |
| *Deinococcus radiodurans* | NC_001263.1 | negative | 3.29 | 7 | 10.73 |
| *Enterococcus faecalis* | NC_017316.1 | positive | 3.36 | 4 | 6.00 |
| *Escherichia coli* | NC_000913.3 | negative | 4.64 | 7 | 7.61 |
| *Lactobacillus gasseri* | NC_008530.1 | positive | 1.89 | 6 | 16.01 |
| *Rhodobacter sphaeroides* | NZ_AKVW01000001.1 | negative | 4.60 | 3 | 3.29 |
| *Staphylococcus epidermidis* | NC_004461.1 | positive | 2.56 | 5 | 9.85 |
| *Streptococcus mutans* | NC_004350.2 | positive | 2.03 | 5 | 12.42 |

**Fig. 5.**
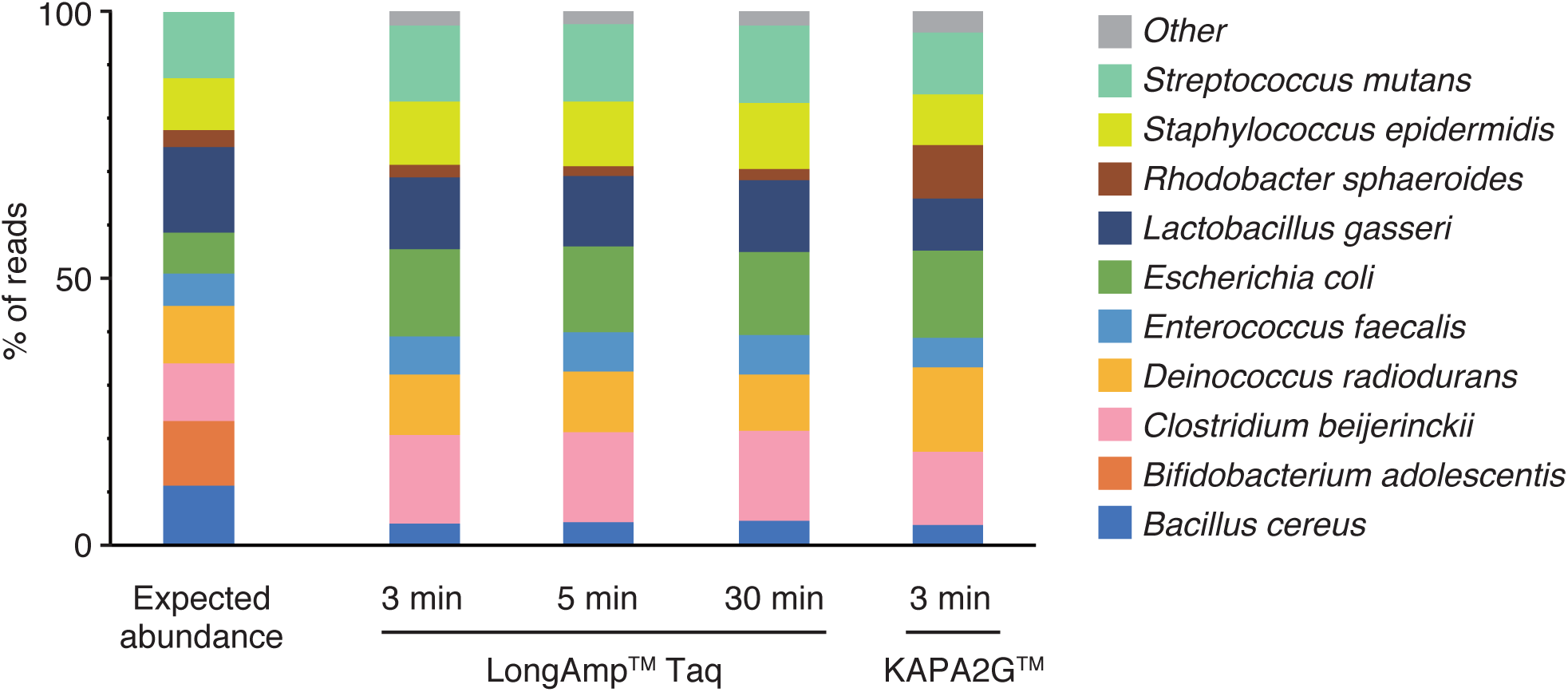
Taxonomic assignment of the mock community consisting of 10 bacterial species. A mixture of DNA from 10 different bacterial species was analyzed by 16S rRNA amplicon sequencing using MinION™. PCR amplification was performed with LongAmp™ Taq or KAPA2G™ polymerase. The samples were analyzed by MinION™ sequencing with different runtime conditions and the percentage of reads mapping to the 10 bacterial species is shown. Expected abundance of individual taxa is based on the genome size and copy number of 16S rRNA genes.

We also evaluated the potential of another DNA polymerase and PCR amplification protocol for bacterial species identification by MinION™ sequencing. KAPA2G™ Robust DNA Polymerase has a significantly faster extension rate than the standard wild-type Taq, enabling shorter reaction times (approximately 100 minutes with LongAmp™ Taq versus 45 minutes with KAPA2G™) for amplifying 16S rRNA genes from the mock bacterial community. The rapid amplification protocol with KAPA2G™ did not impact the overall taxonomy assignment results of 16S rRNA gene sequence reads generated by MinION™ (Fig. 5 and Table 3). Three-minute sequencing of KAPA2G™-amplified 16S rRNA libraries identified nine bacterial species from the mock community. *B. adolescentis* was not detected as was the case with LongAmp™ Taq polymerase.

**Table 3.**
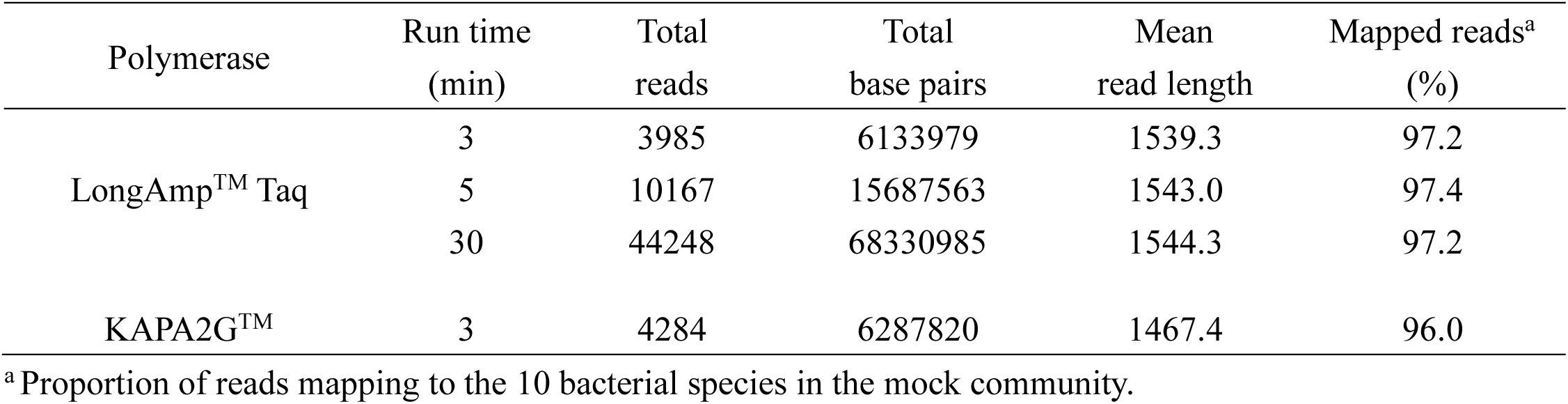
MinION™ sequencing of a bacterial DNA mock community.

### Evaluation of sample preparation methods for accurate bacterial identification via MinION™ sequencing

Given the successful identification of a broad range of bacterial species from mixed DNA samples, we further tested the utility of direct 16S rRNA gene amplification and MinION™ sequencing on a mixture of whole bacterial cells with intact cell walls (Table 4 and Fig. 6). We assessed the effects of DNA extraction procedures on MinION™ sequencing results. Bacterial cell pellets comprising 10 different species were suspended in PBS and divided into three aliquots. The first aliquot remained untreated (designated as “Direct”) and the second was subjected to bead-beating for mechanical cell disruption (“Processed”). Bacterial DNA purified from the third aliquot served as a reference for comparison (“Purified”). Regardless of the extraction procedures, all bacterial species except for *B. adolescentis* were correctly identified and PERMANOVA did not indicate a significant effect for sample preparation methods on community composition (*P* = 0.33). Although not statistically significant, similarity indices may imply that species abundance differed across the three groups (Morisita indices: [Direct:Processed] = 0.66, [Direct:Purified] = 0.65, [Processed:Purified] = 0.87). The relative abundance of *E. coli* was especially high in the 16S rRNA sequencing library amplified directly from the untreated cell suspension and impaired sensitivity was found for the detection of several types of bacteria (Fig. 6A). Mechanical disruption of bacteria by bead-beating can improve results, as it showed patterns of bacterial composition that were more similar to the reference group (Fig. 6B and 6C).

**Table 4.** MinION™ sequencing of a whole cell mock bacterial community.

| Sample | Total reads | Total base pairs | Mean read length | Mapped reads<sup>a</sup> (%) |
| --- | --- | --- | --- | --- |
| Direct | 3868 | 5622803 | 1453.7 | 93.5 |
| Processed (bead-beating) | 3608 | 5500056 | 1524.4 | 97.1 |
| Purified DNA | 3965 | 6035602 | 1522.2 | 96.9 |

<sup>a</sup> Proportion of reads mapping to the 10 bacterial species in the mock community.

**Fig. 6.**
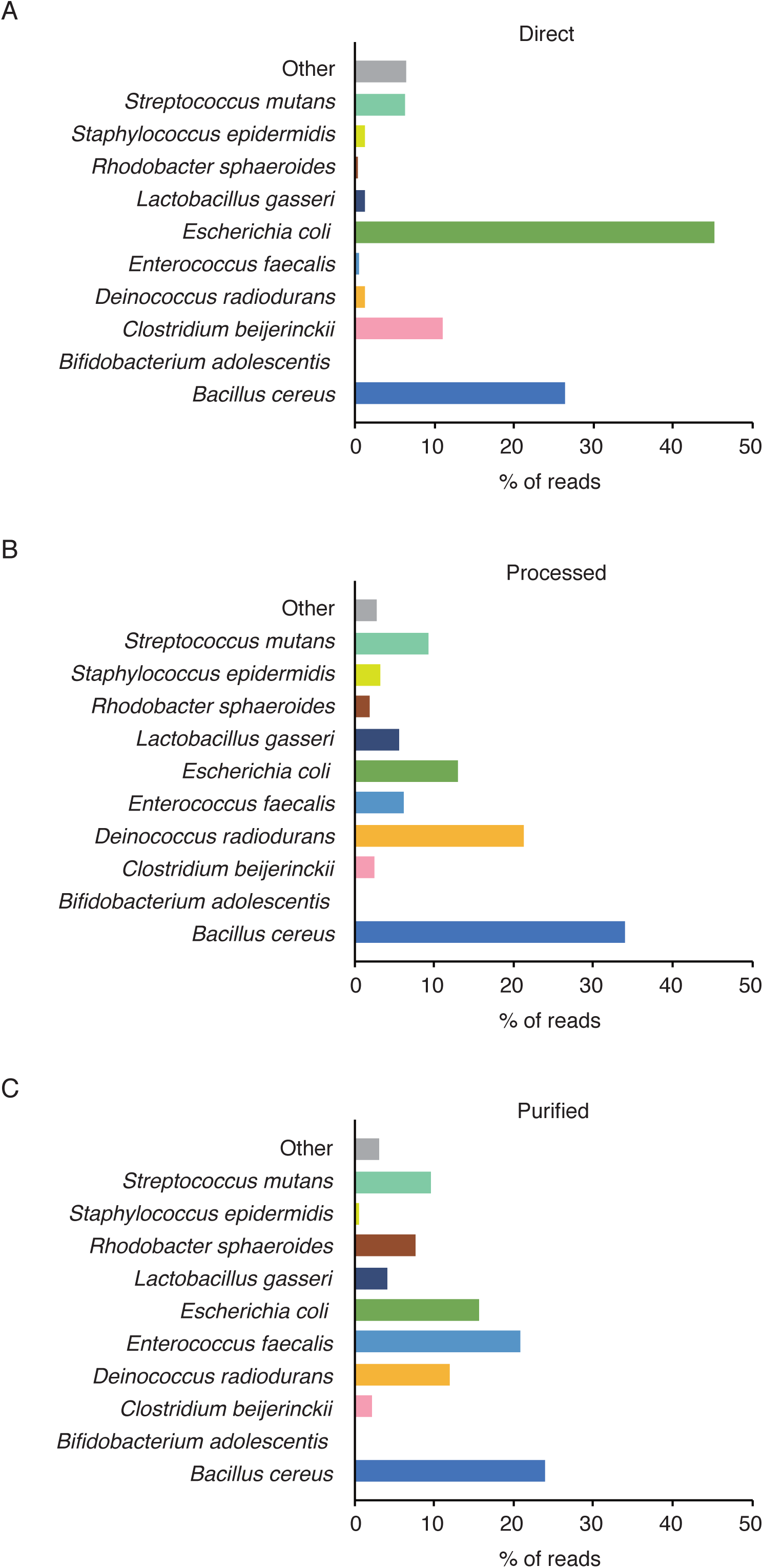
Evaluation of sample processing methods for bacterial composition analysis. (A–C) Whole cell mixtures of 10 bacterial species were (A; direct) left untreated or (B; processed) disrupted by bead-beating for facilitating DNA release (C; purified). Bacterial DNA isolated from the mixed cell suspensions was also used. The samples were subjected to 16S rRNA amplicon sequencing using MinION™ and the percentage of reads mapping to the 10 bacterial species is shown.

## Discussion

Currently, identification of clinically relevant bacteria largely relies on culture-based techniques. However, culture-dependent methods are time intensive and potentially lead to delayed or even incorrect diagnoses [29]. Metagenomic sequencing analysis provides an alternative approach for identifying bacterial pathogens in clinical specimens [5-7]. As previously reported, we developed a sequencing method and bioinformatics pipeline for 16S rRNA gene amplicon sequencing and analysis utilizing the nanopore sequencer MinION™ [15]. Although the system offers faster turnaround time than other NGS platforms, purification of bacterial DNA from samples, which typically takes around 1-2 hours, remains a rate-limiting step in the workflow. Moreover, the bacterial DNA purification requires a multi-step procedure including cell lysis, separation from contaminants, washing, and elution of purified material. These processes are not only time-consuming and laborious but can potentially increase the risk of introducing sample mix-ups and cross contamination. To further facilitate the process of sample preparation, we attempted to amplify 16S rRNA genes directly from bacterial cell suspensions without DNA purification [30, 31]. Three minutes of sequencing run time generated a sufficient number of reads for taxonomic assignment and we achieved successful identification of bacterial species with a total analysis time of less than 2 hours. The direct PCR approach revealed that differences in cell wall morphology (gram status) significantly affected amplification efficiency and sequencing results. For example, *S. aureus*, a gram-positive bacteria with thick cell walls, was more resistant to heat lysis for DNA extraction and yielded less PCR product compared with that of *E. coli*. A mechanical disruption method such as bead-beating was useful in minimizing sample preparation bias. Indeed, samples processed by bead-beating prior to PCR amplification exhibited a better representation of the mock bacterial composition. The differential susceptibility to cell lysis among bacterial species can affect 16S rRNA gene amplification and may introduce a bias in the relative abundance of bacterial species in the community. Thus, mechanical cell disruption preceding direct PCR amplification was indispensable for obtaining an accurate representation of the sample bacterial composition.

We used new primer sets from Oxford Nanopore Technologies that are optimized for rapid 16S rRNA gene sequencing on the MinION™ platform. These universal primers are designed to amplify the near full-length sequence of bacterial 16S rRNA genes. The specificity and sensitivity of 16S rRNA gene amplification with these primers were not substantially affected even when human DNA contaminants outweighed bacterial DNA. Using these primer sets and the updated sequencing protocols, we performed a metagenomic analysis of the pre-characterized bacterial community consisting of 10 different species. Although the universal primers are expected to bind to regions that are highly conserved among bacterial species, we did not detect *B. adolescentis* in the mock community. The 27F forward primer used in this study has three base pair mismatches against *Bifidobacterium* (27F: AGAGTTTGATCMTGGCTCAG; priming site in *B. adolescentis:* AG<u>G</u>GTT<u>C</u>GAT<u>T</u>CTGGCTCA; mismatched bases are underlined) [32]. We speculate that these sequence mismatches lead to poor amplification of *Bifidobacterium* 16S rRNA gene, resulting in the absence of these bacteria in the sequence data. Consistent with our results, it has been reported that the 27F primer has a bias toward underrepresentation of *Bifidobacterium* and other bacterial taxa in microbiome analysis, which is caused by nucleotide variations even in the phylogenetically highly conserved regions of 16S rRNA genes [32-34]. As shown here and in previous publications, it should be noted that universal primers (e.g., 27F primer) commonly used for metagenomic analyses have a limitation related to amplification bias; thus, modifications of primer sequences are required to avoid preferential detection of particular taxa and to cover a broad range of bacterial species [35].

Our study has some limitations. First, the direct PCR approach has been tested only for pure culture of bacteria and a mock community of pre-characterized species. Successful amplification of 16S rRNA genes will greatly depend on the types of biological samples. More extensive studies are required to establish a reliable method for rapid bacterial identification, and future work will focus on optimizing and validating our direct PCR strategy on patient-derived clinical samples. Another issue is that some bacteria share high sequence identity. In this study, *Shigella flexneri* was additionally detected from a pure culture of *E. coli*. Thus, the 16S rRNA gene sequencing has poor discriminatory power to separate closely related species [36, 37]. The sequence analysis targeting additional genetic markers such as 23S rRNA genes may provide better resolution [38].

In conclusion, direct amplification of 16S rRNA genes from crude bacterial extracts can further accelerate sample processing for MinION™ sequencing. Direct 16S rRNA gene amplification combined with MinION™ sequencing provides an attractive option for accelerating pathogen detection. Further optimization and establishment of the relatively simple workflow for rapid bacterial identification via MinION™ sequencing would reduce the turnaround time from sample to result and provide a reliable method that would be applicable to the clinical settings.

## Acknowledgements

This work was supported by grants from the Japanese Society of Anesthesiologists (JSA) Pitch Contest 2017 to SK, JSPS KAKENHI grant number JP16K10975 to YM, MEXT-Supported Program for the Strategic Research Foundation at Private Universities to SN, and Japan Agency for Medical Research and Development (AMED, JP17fm0108023) to TIm. We thank Dr. Koichiro Higasa (Kansai Medical University) for assistance with statistical analysis.

## Author contributions

SK, YM, SN, KK, TIm, and KH designed the study. SK, YM, SM, HT, and TIw conducted the experiments. SK, YM, SN, and KH analyzed and interpreted the data. SK and YM wrote the manuscript.

## Abbreviations

CFU: colony forming unit
PCR: polymerase chain reaction
rRNA: ribosomal RNA

## References

1. Shorr AF, Micek ST, Welch EC, Doherty JA, Reichley RM & Kollef MH (2011) Inappropriate antibiotic therapy in Gram-negative sepsis increases hospital length of stay, Crit Care Med 39: 46–51

2. Puskarich MA, Trzeciak S, Shapiro NI, Arnold RC, Horton JM, Studnek JR, Kline JA, Jones AE & Emergency Medicine Shock Research N (2011) Association between timing of antibiotic administration and mortality from septic shock in patients treated with a quantitative resuscitation protocol, Crit Care Med 39: 2066–2071

3. Fredricks DN & Relman DA (1996) Sequence-based identification of microbial pathogens: a reconsideration of Koch’s postulates, Clin Microbiol Rev 9: 18–33

4. Didelot X, Bowden R, Wilson DJ, Peto TEA & Crook DW (2012) Transforming clinical microbiology with bacterial genome sequencing, Nat Rev Genet 13: 601–612

5. Srinivasan R, Karaoz U, Volegova M, MacKichan J, Kato-Maeda M, Miller S, Nadarajan R, Brodie EL & Lynch SV (2015) Use of 16S rRNA gene for identification of a broad range of clinically relevant bacterial pathogens, PLoS One 10: e0117617

6. Clarridge JE, 3rd (2004) Impact of 16S rRNA gene sequence analysis for identification of bacteria on clinical microbiology and infectious diseases, Clin Microbiol Rev 17: 840–862

7. Kuczynski J, Lauber CL, Walters WA, Parfrey LW, Clemente JC, Gevers D & Knight R (2011) Experimental and analytical tools for studying the human microbiome, Nat Rev Genet 13: 47–58

8. Leggett RM & Clark MD (2017) A world of opportunities with nanopore sequencing, J Exp Bot 68: 5419–5429

9. Jain M, Olsen HE, Paten B & Akeson M (2016) The Oxford Nanopore MinION: delivery of nanopore sequencing to the genomics community, Genome Biol 17: 239

10. Quick J, Ashton P, Calus S, Chatt C, Gossain S, Hawker J, Nair S, Neal K, Nye K, Peters T, De Pinna E, Robinson E, Struthers K, Webber M, Catto A, Dallman TJ, Hawkey P & Loman NJ (2015) Rapid draft sequencing and real-time nanopore sequencing in a hospital outbreak of Salmonella, Genome Biol 16: 114

11. Sanderson ND, Street TL, Foster D, Swann J, Atkins BL, Brent AJ, McNally MA, Oakley S, Taylor A, Peto TEA, Crook D & Eyre DW (2017) Real-time analysis of nanopore-based metagenomic sequencing from orthopaedic device infection, bioRxiv doi: https://doi.org/10.1101/220616

12. Leggett RM, Alcon-Giner C, Heavens D, Caim S, Brook TC, Kujawska M, Hoyles L, Clarke P, Hall L & Clark MD (2017) Rapid MinION metagenomic profiling of the preterm infant gut microbiota to aid in pathogen diagnostics, bioRxiv doi: https://doi.org/10.1101/180406

13. Schmidt K, Mwaigwisya S, Crossman LC, Doumith M, Munroe D, Pires C, Khan AM, Woodford N, Saunders NJ, Wain J, O’Grady J & Livermore DM (2017) Identification of bacterial pathogens and antimicrobial resistance directly from clinical urines by nanopore-based metagenomic sequencing, J Antimicrob Chemother 72: 104–114

14. Benitez-Paez A, Portune KJ & Sanz Y (2016) Species-level resolution of 16S rRNA gene amplicons sequenced through the MinION portable nanopore sequencer, GigaScience 5: 4

15. Mitsuhashi S, Kryukov K, Nakagawa S, Takeuchi JS, Shiraishi Y, Asano K & Imanishi T (2017) A portable system for rapid bacterial composition analysis using a nanopore-based sequencer and laptop computer, Sci Rep 7: 5657

16. Shin H, Lee E, Shin J, Ko SR, Oh HS, Ahn CY, Oh HM, Cho BK & Cho S (2018) Elucidation of the bacterial communities associated with the harmful microalgae Alexandrium tamarense and Cochlodinium polykrikoides using nanopore sequencing, Sci Rep 8: 5323

17. Ma X, Stachler E & Bibby K (2017) Evaluation of Oxford Nanopore MinION Sequencing for 16S rRNA Microbiome Characterization, bioRxiv doi: https://doi.org/10.1101/099960

18. Miller JH (1972) Determination of viable cell counts: bacterial growth curves in Experiments in Molecular Genetics, pp. 31–36, Cold Spring Harbor Laboratory Press, Cold Spring Harbor, NY.

19. Jiang H, Dong H, Zhang G, Yu B, Chapman LR & Fields MW (2006) Microbial diversity in water and sediment of Lake Chaka, an athalassohaline lake in northwestern China, Appl Environ Microbiol 72: 3832–3845

20. Eden PA, Schmidt TM, Blakemore RP & Pace NR (1991) Phylogenetic analysis of Aquaspirillum magnetotacticum using polymerase chain reaction-amplified 16S rRNA-specific DNA, Int J Syst Bacteriol 41: 324–325

21. Frith MC (2011) A new repeat-masking method enables specific detection of homologous sequences, Nucleic Acids Res 39: e23

22. Li H (2018) Minimap2: pairwise alignment for nucleotide sequences, Bioinformatics 34: 3094–3100

23. Federhen S (2012) The NCBI Taxonomy database, Nucleic Acids Res 40: D136–143

24. Ondov BD, Bergman NH & Phillippy AM (2011) Interactive metagenomic visualization in a Web browser, BMC Bioinformatics 12: 385

25. Morisita M (1959) Measuring of interspecific association and similarity between communities, Mem Fac Sci Kyushu Univ Series E 3: 65–80

26. Oksanen J, Blanchet FG, Friendly M, Kindt R, Legendre P, McGlinn D, Minchin PR, O’Hara RB, Simpson GL, Solymos P, Stevens MHH, Szoecs E & Wagner H (2017) vegan: Community Ecology Package. R package version 2.4–5. https://CRAN.R-project.org/package=vegan

27. Fukushima M, Kakinuma K & Kawaguchi R (2002) Phylogenetic analysis of Salmonella, Shigella, and Escherichia coli strains on the basis of the gyrB gene sequence, J Clin Microbiol 40: 2779–2785

28. Devanga Ragupathi NK, Muthuirulandi Sethuvel DP, Inbanathan FY & Veeraraghavan B (2018) Accurate differentiation of Escherichia coli and Shigella serogroups: challenges and strategies, New Microbes New Infect 21: 58–62

29. Clarridge JE, 3rd, Raich TJ, Sjosted A, Sandstrom G, Darouiche RO, Shawar RM, Georghiou PR, Osting C & Vo L (1996) Characterization of two unusual clinically significant Francisella strains, J Clin Microbiol 34: 1995–2000

30. Videvall E, Strandh M, Engelbrecht A, Cloete S & Cornwallis CK (2017) Direct PCR Offers a Fast and Reliable Alternative to Conventional DNA Isolation Methods for Gut Microbiomes, mSystems 2: e00132–00117

31. Flores GE, Henley JB & Fierer N (2012) A direct PCR approach to accelerate analyses of human-associated microbial communities, PLoS One 7: e44563

32. Walker AW, Martin JC, Scott P, Parkhill J, Flint HJ & Scott KP (2015) 16S rRNA gene-based profiling of the human infant gut microbiota is strongly influenced by sample processing and PCR primer choice, Microbiome 3: 26

33. Frank JA, Reich CI, Sharma S, Weisbaum JS, Wilson BA & Olsen GJ (2008) Critical evaluation of two primers commonly used for amplification of bacterial 16S rRNA genes, Appl Environ Microbiol 74: 2461–2470

34. Farris MH & Olson JB (2007) Detection of Actinobacteria cultivated from environmental samples reveals bias in universal primers, Lett Appl Microbiol 45: 376–381

35. von Wintzingerode F, Gobel UB & Stackebrandt E (1997) Determination of microbial diversity in environmental samples: pitfalls of PCR-based rRNA analysis, FEMS Microbiol Rev 21: 213–229

36. Jenkins C, Ling CL, Ciesielczuk HL, Lockwood J, Hopkins S, McHugh TD, Gillespie SH & Kibbler CC (2012) Detection and identification of bacteria in clinical samples by 16S rRNA gene sequencing: comparison of two different approaches in clinical practice, J Med Microbiol 61: 483–488

37. Chatellier S, Mugnier N, Allard F, Bonnaud B, Collin V, van Belkum A, Veyrieras JB & Emler S (2014) Comparison of two approaches for the classification of 16S rRNA gene sequences, J Med Microbiol 63: 1311–1315

38. Kerkhof LJ, Dillon KP, Haggblom MM & McGuinness LR (2017) Profiling bacterial communities by MinION sequencing of ribosomal operons, Microbiome 5: 116

